# MultiAlloDriver: a multi-model method to predict and identify cancer driver mutations

**DOI:** 10.1101/2025.06.25.661496

**Authors:** Wanyao Zhou, Yuchengze Song, Jixiao He, Mingyu Li, Jian Zhang, Shaoyong Lu

## Abstract

A minority of driver mutations in cancer significantly alter protein structure and key functionalities, thereby driving cancer progression. Consequently, the prediction and identification of driver mutations hold critical implications for targeted cancer therapy. This study introduces MultiAlloDriver, a novel multi-modal machine learning model based on an attention mechanism, which for the first time, incorporates protein surface information. By integrating information from three dimensions-protein sequence, structure, and surface-the model achieves high-accuracy driver target prediction with an accuracy exceeding 93%. Notably, we utilized this tool to predict and identify the driver effect of the F90S mutation in the PTEN tumor suppressor gene, uncovering mutations associated with cancer signaling mechanisms. Overall, MultiAlloDriver contributes to elucidating the underlying mechanisms of cancer development and progression, providing a robust framework for the identification of driver targets.

## Introduction

Cancer is a disease caused by the cumulative effects of mutations^1–3^. During its development, a small subset of mutations that significantly alter protein structure, phenotype, or function, thereby driving carcinogenesis, are termed driver mutations^4^, while the majority of mutations with minimal impact on protein structure or function are classified as passenger mutations. Numerous studies have demonstrated that driver mutations are closely associated with critical processes in cancer progression, including signal transduction, aberrant proliferation, metastasis, and drug resistance^5–7^. Consequently, a pivotal scientific challenge in cancer-targeted therapy lies in the accurate prediction and identification of driver mutations. High-precision detection of mutation types not only elucidates potential mechanisms underlying cancer progression but also facilitates the discovery of key therapeutic targets and guides the development of targeted anticancer drugs.

Mutations in protein-coding regions can alter amino acid sequences, leading to modified local folding patterns, structural perturbations (local or global), and ultimately functional changes that disrupt interactions with upstream/downstream signaling molecules^8^. Based on their topological and spatial relationships with functional domains, mutation sites can be categorized as orthosteric or allosteric^9–11^. Orthosteric sites reside within functional regions, where mutations directly interfere with domain structure and function, thereby impacting molecular interactions. In contrast, allosteric sites are distal to functional regions. Emerging evidence suggests that such mutations may indirectly modulate functional domains through long-range structural rearrangements (e.g., altered hydrogen bonding networks or conformational dynamics)^12–14^. However, due to mechanistic complexity and limitations of traditional methodologies, allosteric driver mutations have not been as extensively studied as their orthosteric counterparts. Thus, the prediction and characterization of allosteric driver mutations hold significant potential for uncovering novel therapeutic targets in cancer.

Protein surface information, a high-level representation derived from boundary atoms forming a mesh-like structure, encodes critical physicochemical properties such as geometric configuration, charge distribution, hydrophobicity, and hydrogen-bonding capacity. Studies indicate that proteins with highly similar sequences or structures may exhibit divergent molecular binding affinities due to surface variations, whereas surface similarity often correlates with conserved biological activity^15,16^. Compared to sequence or structural data, surface features more directly govern protein interaction capabilities, particularly for functional domains^17^. For many driver mutations, structural destabilization caused by residue substitutions frequently coincides with localized surface alterations (e.g., hydrophobicity shifts or charge redistribution), which may directly influence folding pathways and secondary structure formation. KRAS, a GTPase belonging to the RAS family, serves as a critical molecular switch in cellular signal transduction, particularly regulating pathways governing cell growth, proliferation, and survival^18^. In its GTP-bound state, KRAS adopts an active conformation to propagate downstream signals, whereas hydrolysis of GTP to GDP transitions it to an inactive state—a cycle tightly regulated under physiological conditions^18,19^. KRAS mutations are strongly associated with malignancies such as pancreatic, colorectal, and non-small cell lung cancers^20,21^. Among these, the G12V variant represents one of the most oncogenic mutations. This substitution introduces a hydrophobic residue that impairs GTPase-mediated hydrolysis, locking KRAS in a constitutively active GTP-bound state^22^. Clinically, G12V has been linked to aggressive tumor phenotypes and poor prognosis^23^. Structural analyses of wild-type versus mutant KRAS (Fig. 1) reveal that the G12V missense mutation induces significant local backbone rearrangements, reshaping the surface topology of the GTP-binding pocket^24^. These alterations correlate with diminished GTPase activity, underscoring the interplay between mutational effects and surface feature remodeling. Anaplastic lymphoma kinase (ALK) is a member of the insulin receptor protein-tyrosine kinase superfamily. Under physiological conditions, ALK is activated by phosphorylation at Y1258 and regulates cell proliferation, differentiation, and survival through downstream signaling pathways (e.g., PI3K/AKT, MAPK/ERK, JAK/STAT), primarily functioning during central nervous system development. Aberrant ALK activation has been implicated in diverse malignancies, including anaplastic large-cell lymphoma, lung cancer, and neuroblastoma^25–27^. The F1174L mutation is a well-characterized pathogenic allosteric driver mutation in ALK. Structural studies demonstrate that substitution of phenylalanine with leucine at position 1174 promotes tighter interactions between the αC-helix and proximal A-loop helix, while inducing distal A-loop distortion^28^. This conformational rearrangement exposes the P+1 pocket, leading to ligand-independent constitutive kinase activation even in the absence of Y1258 phosphorylation. Further analysis of the protein structure revealed that the F1174L mutation induces significant alterations in the geometry of adjacent surface pockets. The DEAD-box RNA helicase DDX3X belongs to the Superfamily 2 (SF2) of RNA helicases and contains two tandem RecA-like domains (D1 and D2)^29^. It catalyzes ATP hydrolysis and remodels RNA secondary structure through several conserved helicase motifs. DDX3X plays critical roles in various cellular processes, including the regulation of translation of mRNAs with complex 5′ untranslated regions (UTRs). Loss of DDX3X function has been implicated in several cancers, including medulloblastoma^30^. Among the disease-associated mutations, D354V occurs at the interface between the D1 and D2 domains. This mutation replaces a hydrophilic, negatively charged aspartic acid (D) with a more hydrophobic valine (V), thereby disrupting electrostatic interactions and increasing hydrophobicity at the domain interface. Crystallographic analysis indicates that while the D354V mutation does not significantly alter the individual structures of D1 or D2, it does induce a substantial rearrangement of their relative orientation, with D2 rotated approximately 180° compared to the wild type. This conformational shift results in an “open but dynamic” structure, which likely contributes to crystallization stability^31^. More importantly, although D354V does not directly impair canonical ATP-binding motifs, it affects the N-terminal ATP-binding loop (ABL, residues 135–168), thereby indirectly reducing RNA-stimulated ATPase activity. The mutant retains catalytic ATP hydrolysis capacity; however, it exhibits only mild defects in dsRNA binding and ATPase stimulation in the presence of double-stranded RNA. These findings suggest that D354V compromises the protein’s responsiveness to specific RNA structures, which may underlie its pathogenic effects. The protein tyrosine kinase JAK2 is a key signaling molecule downstream of a wide range of cytokine receptors, including those for erythropoietin, growth hormone, interferon-γ, and interleukins such as IL-3 and IL-5. It functions via the JAK-STAT signaling pathway to regulate hematopoietic cell proliferation, differentiation, and immune responses^32^. JAK2 is composed of several conserved domains: an N-terminal FERM domain responsible for receptor binding, a SH2-like domain with unclear function, a pseudokinase domain (JH2), and a C-terminal catalytic tyrosine kinase domain (JH1)^33^. Mutations in the JH2 domain of JAK2 are strongly associated with myeloproliferative neoplasms (MPNs), which are clonal hematopoietic disorders characterized by excessive proliferation of myeloid lineages. The V617F mutation in JH2 is the most common pathogenic alteration and may lead to the development of polycythemia vera, essential thrombocythemia, and primary myelofibrosis. Structurally, V617F replaces a small, nonpolar valine with a bulky aromatic phenylalanine, enhancing hydrophobic interactions and rigidifying the adjacent αC-helix within the N-lobe of the JH2 domain. This conformational stabilization promotes the stimulatory interaction of JH2 with JH1, facilitating constitutive trans-phosphorylation of the activation loop in JH1 independent of cytokine stimulation^34^. Computational prediction using FPocket confirmed substantial differences in both pocket volume and druggability scores, underscoring a direct correlation between the mutation and structural remodeling of surface pockets.

**Figure 1:**
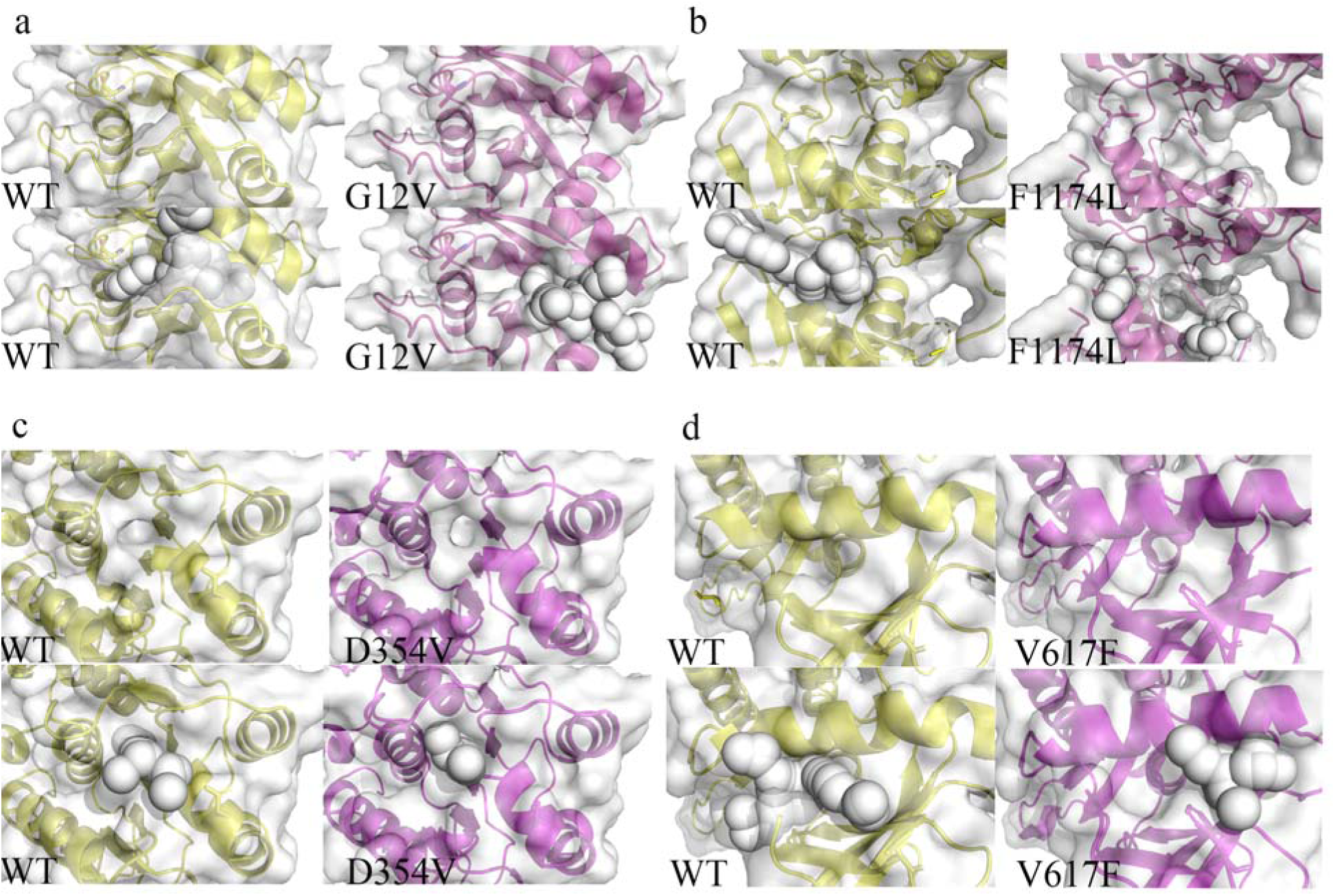
Protein surface change before and after mutation, the original one and the one after using FPocket to represent the structure. **a.** G12V mutation in KRAS. **b.**F1174L in ALK. c.D354V in DDX3X. d.V617F in JAK2.

**Figure 2.**
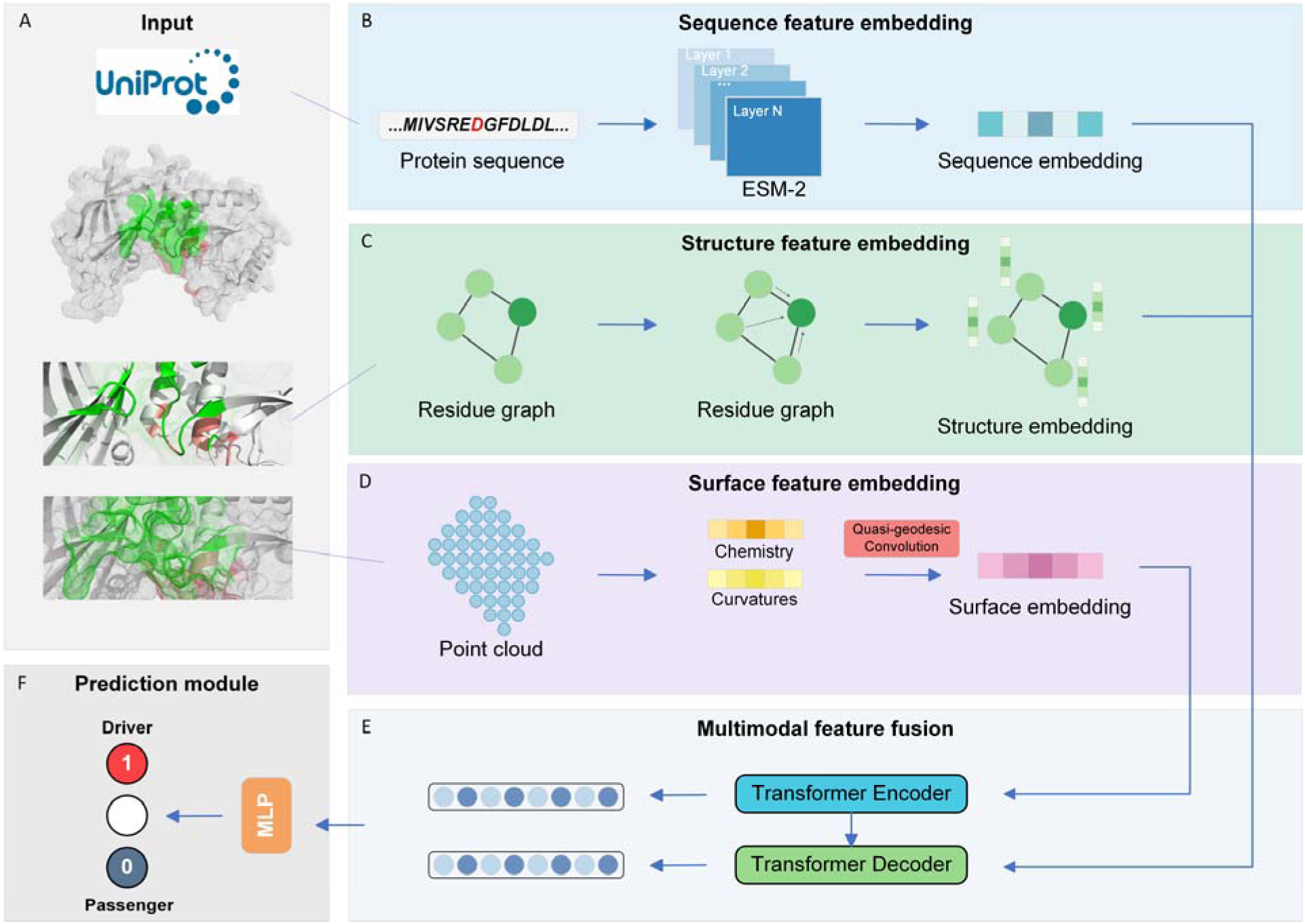
The workflow of MultiAlloDriver.

**Figure 3.**
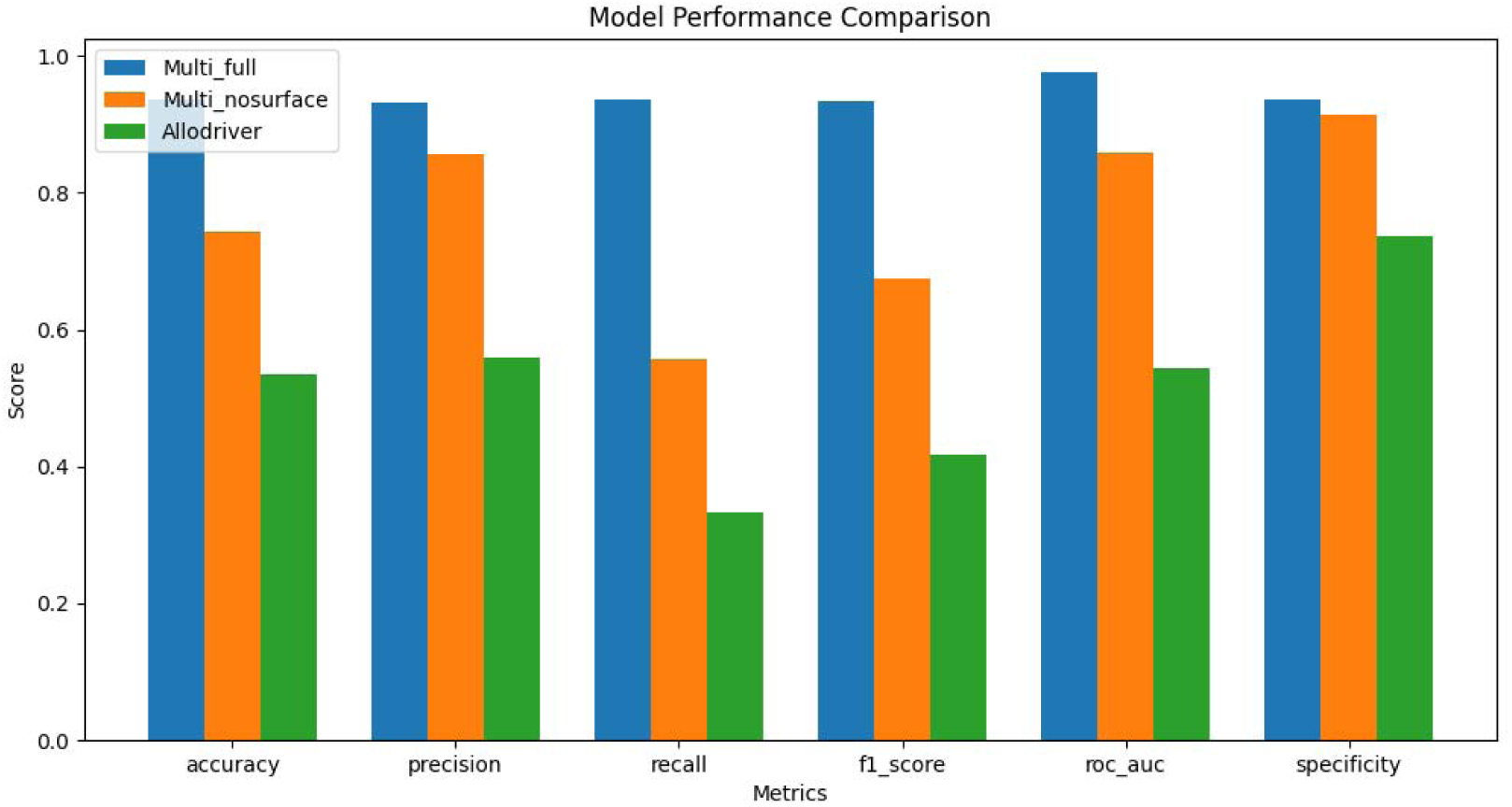
Bar plots comparing performance metrics between the full MultiAlloDriver model (blue) and the surface-ablated model (yellow). Key observations: accuracy (−21%), recall (−41%), and AUC (−12%) exhibited substantial reductions upon surface feature removal.

**Figure 4.**
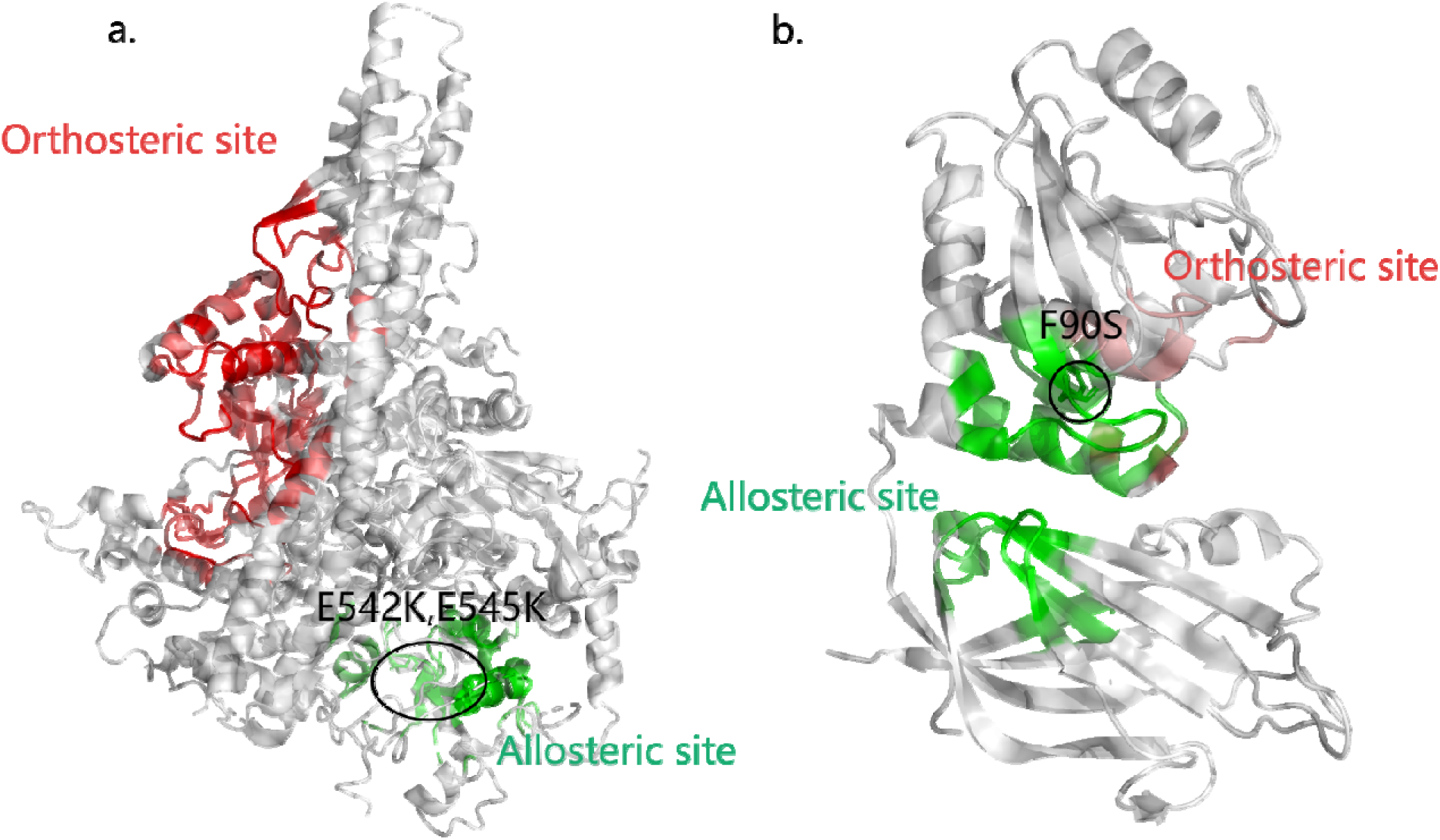
Examples of MultiAlloDriver. **a.**The driver mutation E542K, E545K is found at the PI3Kα allosteric site (PDB ID: 4OVV). The protein structure is exhibited in both cartoon and transparent surface modes with allosteric site coloured in green and orthosteric site in red. **b.** The driver mutation F90S is found at the PTEN allosteric site (PDB ID:1D5R). The protein structure is exhibited in both cartoon and transparent surface modes with allosteric site coloured in green and orthosteric site in red.

Correlatively, James W. McCormick et al. (2021) revealed that the allosteric H124Q mutation in Dihydrofolate Reductase markedly increased E. coli photoresponse, with synergistic mutations distributed across the protein surface amplifying allosteric effects^35^. Similarly, Nikita R. Dsouza et al. (2024) demonstrated that mutations in Granzyme H and IGF1R statistically altered surface charge distributions, explaining crystallographic disparities between wild-type and mutant proteins^16^. These findings underscore the critical role of surface properties in mediating mutation-induced functional changes, highlighting their importance for allosteric driver site identification.

While surface information has been widely applied in drug design and molecular docking^36^, its inherent complexity, high-dimensional representation, and technical challenges in feature extraction historically limited its utilization. Recent advances in machine learning, however, have enabled deep mining of surface features. The Masif model (2020)^37^, employing geodesic convolutional neural networks to process surface meshes, demonstrated robust performance in molecular docking but suffered from non-end-to-end processing and high memory demands. Subsequently, the dMasif framework addressed these limitations by representing surfaces as atomic point clouds, achieving enhanced computational efficiency and performance^38^.Despite advances facilitated by AlphaFold and machine learning, current models for predicting allosteric driver sites predominantly rely on sequence and structural features^39–42^ while neglecting surface information—a key determinant of molecular interactions, especially for surface-localized allosteric mutations. To address this gap, we propose MultiAlloDriver, a multimodal deep learning framework integrating sequence, structure, and surface features based on our past research^43,44^. Our approach employs: (1) ESM-2 for extracting long-range sequence dependencies, (2) GVP-GNNs for structural graph representations, and (3) dMasif-based atomic point clouds for surface feature encoding. A transformer-based attention mechanism aligns and fuses these multimodal features. MultiAlloDriver achieved 94% accuracy and precision on test datasets. Furthermore, we validated its utility by identifying the F90S allosteric driver mutation in the tumor suppressor PTEN, elucidating its impact on dephosphorylation activity. As the first model incorporating surface information for mutation effect prediction, MultiAlloDriver unveils novel cancer mechanisms and provides a powerful tool for guiding targeted therapy development.Materials and Methods.

### Performance

We randomly divided the 17,130 allosteric driver and passenger mutations into training, validation, and test sets in an 8:1:1 ratio (13,704 for training, 1,713 for validation, and 1,713 for testing).

Using these data, we built multimodal input features to train and assess a driver-mutation recognition model. The resulting deep network fuses primary protein sequence, 3-D backbone geometry, and surface chemo-geometric properties to discriminate drivers from passengers. After training, the model outputs continuous probabilities on previously unseen data; following the literature, scores above 0.50 are labeled as drivers, otherwise passengers.

On the validation set the model achieved an accuracy of 0.9348, and the overall accuracy reached 0.9365. The confusion-matrix metrics show balanced precision (0.9314) and recall (0.9371), yielding an F-score of 0.9342—indicating a high detection rate for driver mutations while controlling false positives. Crucially, the threshold-independent ROC-AUC reached 0.97, far exceeding the 0.80 benchmark for “excellent” performance and demonstrating an ideal trade-off between true- and false-positive rates across all thresholds. Together, these results place our framework at the forefront of current methods in both accuracy and robustness, offering a reliable tool for large-scale functional annotation of cancer variants and for clinical precision-medicine screening.

### Surface Feature Ablation Analysis

To evaluate the contribution of surface features to model performance and validate their importance in protein feature extraction, we conducted ablation experiments (Figure n) comparing the complete MultiAlloDriver model with a degraded version lacking surface information inputs. Performance metrics were systematically compared between the full model (integrating sequence, structure, and surface features) and the ablated model (using only sequence and structural data). Results demonstrated significant declines across all metrics upon surface feature removal: accuracy decreased by 21%, AUC dropped by 12%, and recall exhibited the most pronounced reduction (41%). These findings confirm that surface information effectively enhances protein feature representation and substantially improves predictive capability for cancer-related allosteric driver mutations.

### Case Studies

#### 1. PI3K**α** (Phosphoinositide 3-Kinase Alpha)

As a critical member of the class I PI3K family, PI3Kα functions as a heterodimer comprising a catalytic subunit (p110α) and a regulatory subunit (p85α/β). The p110α subunit catalyzes the phosphorylation of phosphatidylinositol 4,5-bisphosphate (PIP2) to phosphatidylinositol 3,4,5-trisphosphate (PIP3), while the p85 subunit modulates enzymatic activity and mediates interactions with upstream signaling molecules (e.g., receptor tyrosine kinases). PI3Kα plays a central role in the PI3K/AKT/mTOR pathway, regulating cell growth, proliferation, survival, and metabolism. Dysregulated PI3Kα activation is strongly associated with malignancies such as breast, colorectal, and endometrial cancers. MultiAlloDriver identified the E542K and E545K mutations within the helical domain of p110α as high-confidence allosteric driver sites (prediction probabilities: 99.74% and 99.56%, respectively). These mutations localize to a key allosteric pocket, where they disrupt interfacial salt bridges, relieve autoinhibition, and induce active-site remodeling—mechanisms corroborated by prior studies linking these variants to constitutive PI3Kα activation^14,45^.

#### 2. PTEN (Phosphatase and Tensin Homolog)

PTEN, a pivotal tumor suppressor, antagonizes PI3K signaling by dephosphorylating PIP3 to PIP2, thereby inhibiting the PI3K/AKT/mTOR pathway and regulating apoptosis, proliferation, and migration. Its functional domains include the phosphatase domain, C2 domain (mediating membrane binding), and PDZ-binding motif. MultiAlloDriver predicted the F90S mutation within the phosphatase domain as a potent allosteric driver (probability: 99.86%). Structural mapping revealed that F90S resides in an allosteric network involving Phe90, Glu91, Asp92, His93, Asn94, Pro95, Pro96, Gln97, Gln219, Leu220, Lys221, and Asp252. This mutation induces long-range allosteric effects that impair C2 domain-membrane interactions (Reference), significantly reducing PTEN’s dephosphorylation activity^46^. These results highlight the therapeutic potential of targeting F90S for novel anticancer drug development.

## Methods

### Dataset

The dataset used in this study was compiled by our group in earlier work and was originally collected and constructed systematically by [Author Name] et al. (see Reference [XX])^47^. It integrates driver-mutation information from multiple public databases, including the Cancer Genome Interpreter (CGI)^48^, Integrative OncoGenomics (IntOGen)^49^, Clinical Interpretation of Variants in Cancer (CIViC)^50^, the Database of Curated Mutations (DoCM)^51^, MSK’s OncoKB^52^, and the Precision Medicine Knowledgebase (PMKB)^53^. After rigorous standardization and filtering, we obtained 715 driver genes with a total of 17,181 mutation records, among which 8,565 are confirmed driver mutations.

To create a balanced training set, we randomly selected an equal number of passenger mutations from other sources, yielding a balanced dataset of 17,130 samples comprising both driver and passenger mutations.

### Mutimodal structure Surface feature embedding

Protein surfaces are rich in chemical and geometric features that directly reflect interactions between a mutation site and nearby residues^54^. Unlike the costly, non-differentiable pre-computations required by MaSIF, dMaSIF employs a fully end-to-end differentiable network that takes only atom types and 3-D coordinates as input, efficiently reconstructing and encoding the local geometry and chemical environment of the surface^55^. Concretely, for each mutation site we extract an allosteric site by cropping the 512 atoms lying within a ∼7 Å radius. From this region we derive surface descriptors—such as curvature distribution, charge density, and the spatial arrangement of polar groups—and finally generate a fixed-dimensional embedding vector that represents the allosteric site.

dMaSIF treats a protein as an unordered point cloud of atoms. For each atom type, it first builds a 22-dimensional one-hot vector to encode its chemical identity. It then assigns a characteristic influence radius to every atom type and, on that basis, defines a smooth distance field that gives the shortest distance from any point in space to the molecular surface. By solving the iso-distance condition on this field, the method uses numerical optimization to uniformly sample surface points at the target distance, achieving a resolution of about 1 Å and typically producing several thousand to tens of thousands of points—enough to cover the entire exterior and capture fine concave-convex details.

At each sampled point, the algorithm computes the gradient of the distance field to estimate the surface normal. It then constructs two tangent vectors in the plane orthogonal to this normal. Together, the normal and the two orthogonal tangent vectors define a local 3-D coordinate frame. This frame provides invariance to rigid-body motions and local rotations in the downstream network, enabling a robust encoding of protein shape.

For chemical feature extraction, we query the 16 nearest atomic centers around each surface point. The one-hot encodings of these atoms are passed through a lightweight multilayer perceptron (MLP), whose nonlinear mapping yields a fixed-dimensional chemical embedding.

For geometric feature extraction, we combine the curvature information at that point with the directions of the local tangent vectors. Using either local surface fitting or spectral techniques, we derive a geometric vector that captures the degree of concavity/convexity and bending of the surface patch.

The chemical embedding and geometric vector are then concatenated to form a multimodal feature vector for every surface point. This vector simultaneously encodes the chemical environment of nearby atomic clusters and the fine-grained morphology of the molecular surface, supplying downstream deep networks with a comprehensive, compact, and fully differentiable representation—thereby enhancing the prediction of allosteric sites and the functional impact of driver-mutation sites.

To further aggregate the multimodal features of surface points and generate the final embedding, we employ quasi-geodesic convolution. Designed specifically for protein surface– oriented point clouds, this algorithm propagates information along approximate geodesic paths within local neighborhoods, allowing the network to adaptively learn feature representations pertinent to driver-mutation prediction. Unlike conventional Euclidean convolutions, quasi-geodesic convolution is naturally equivariant or invariant to 3-D rotations and translations— meaning that, regardless of how the protein is oriented in space, the resulting convolved features remain consistent.

Concretely, for each sampled surface point the method first constructs an approximate geodesic neighborhood using the local tangent vector field. Within this neighborhood it defines a set of learnable filter kernels, which aggregate features through weighted summation. Stacking multiple such convolutional layers enables the network to capture progressively larger surface-context information. This process preserves fine-scale geometric curvature and chemical-distribution details while distilling global conformational patterns layer by layer, providing the downstream multimodal network with a highly robust and biologically meaningful representation that markedly enhances the identification and classification of driver-mutation sites.

### Structure feature embedding

In deep learning, a protein’s three-dimensional structure encodes abundant physicochemical information, clearly revealing how each residue interacts with its surrounding amino acids and how signals propagate through the molecule. To mine these structural cues efficiently, a protein is typically abstracted as a residue graph—nodes represent residues (or atoms), while edges link nodes according to spatial proximity or chemical bonds—and a graph neural network (GNN) serves as the core feature extractor.

Through an iterative message-passing scheme, a GNN captures not only local inter-residue dependencies but, when multiple layers are stacked, the protein’s global topology as well. At each layer, a node receives messages from its neighbors; these messages are generated by learnable functions that combine the neighbors’ features with relative spatial encodings. An aggregation operation (such as a weighted sum or normalized mean) pools the messages into a neighborhood representation, which is then fused with the node’s own features to update its embedding. Modern GNN designs commonly incorporate residual connections and layer normalization between layers to ensure stable information flow in deep networks and to reduce over-smoothing.

By jointly leveraging residue sequences, chemical attributes, and geometric coordinates, GNN-based models have delivered strong results across a variety of tasks, including functional prediction, active-site identification, protein–protein interaction classification, and drug-binding affinity estimation^56,57,58^.

To achieve simultaneous invariance to 3-D rotations and translations, we embed a Geometric Vector Perceptron (GVP) into the standard message-passing framework^59^. Designed for biomolecular data, GVP separates each node’s representation into scalar and vector channels. In the message and update functions, scalar and vector features are processed through independent linear projections followed by nonlinear activations. Before producing the new scalar output, the L norm of the incoming vector channel is concatenated to the projected scalar features, preserving rotational invariance. The new vector output is obtained by applying a linear transform directly to the input vector, ensuring equivariance with respect to 3-D rotations.

Through this mechanism, each round of message passing updates chemical and sequence information while precisely capturing local geometric directions and deformation trends, yielding node embeddings that remain biologically meaningful and robust to rigid-body motions.

Our network also adopts a multi-scale aggregation strategy in which skip connections fuse GVP outputs from different depths. By integrating signals that span from atomic to whole-protein scales, the model can detect subtle geometric changes in local allosteric pockets as well as large-scale backbone rearrangements. This comprehensive structural awareness significantly enhances driver-mutation classification performance and supports more insightful biological interpretation.

### Sequence feature embedding

ESM-2 (Evolutionary Scale Modeling 2) is currently the best-performing general-purpose protein language model^60^. Built on a Transformer architecture, it has been self-supervised on large sequence repositories such as UniProt via a masked-language-modeling objective, enabling it to capture evolutionary signals and latent structural information directly from sequences. Beyond predicting structure and function for single proteins, the model supports a wide array of downstream tasks, including active-site identification and discrimination of protein–protein interaction interfaces^61,62^.

In this study we benchmarked four differently sized ESM-2 variants: a 6-layer model with 8 million parameters, a 12-layer model with 35 million parameters, a 30-layer model with 150 million parameters, and a 33-layer model with 650 million parameters. Although performance generally increases with model size, larger models impose greater demands on computation and GPU memory. Balancing accuracy gains against hardware constraints, we ultimately selected the largest variant—ESM2_t33_650M_UR50D, containing 33 layers and roughly 650 million parameters—for our experiments.

In practice, every layer of the model assigns a 1,280-dimensional hidden vector to each amino-acid residue in the sequence. These vectors fuse local sequence context with information about evolutionary conservation and structural propensities. We first retrieve the full amino-acid sequence of the target protein via its UniProt ID and replace the wild-type residue at the mutation site to obtain the mutated sequence. Both sequences are then passed through the pre-loaded ESM2_t33_650M_UR50D model, from which we extract the 1,280-dimensional residue embeddings produced by the deepest—33rd—layer; prior studies indicate that representations from the final layer capture higher-level abstractions capable of distinguishing diverse functional sites.

Finally, we perform global average pooling over the entire sequence of residue vectors to produce a fixed-length sequence feature. This vector not only quantifies the impact of a single-point mutation in the sequence-representation space but also integrates seamlessly with structural and surface-level modalities, providing a rich and efficient descriptor for accurately classifying driver versus passenger mutations.

### Multimodal feature fusion

In practical settings, combining multiple modalities poses a number of challenges. The data distributions and feature scales differ markedly across modalities: sequence embeddings encode linguistic information as high-dimensional dense vectors, whereas surface point-cloud features are sparse geometric–chemical descriptors. Simple concatenation or naïve blending can let one modality dominate while drowning out the others. Moreover, sequence, structural, and surface data are neither spatially nor sequentially synchronized, and without a clear alignment mechanism it is difficult to establish precise correspondences among them.

Multimodal fusion is still hampered by heterogeneous data distributions, mismatched feature scales, and the lack of an explicit alignment mechanism—issues that can cause one modality to dominate and make it difficult to establish precise correspondences across sequence, structure, and surface representations. Against this backdrop, the Transformer architecture offers natural advantages. It can model serialized features and capture long-range dependencies through self-attention, and—by means of cross-attention—build direct links between different feature spaces, enabling deep alignment and complementary integration of information.

We first feed the surface-point features extracted by dMaSIF into a dedicated Transformer encoder. Multi-head self-attention lets the model evaluate similarity and interaction between any two points globally, automatically focusing on critical regions such as surface pockets and charge patterns. Subsequent feed-forward layers, together with residual connections and layer normalization, yield a high-dimensional representation that optimally summarizes global surface geometry. In parallel, we project the ESM-2 sequence embeddings and the GVP-processed graph embeddings through separate linear layers and add positional encodings to preserve both backbone order and spatial relationships. Each is then refined by an additional self-attention layer, which amplifies long-range co-evolutionary signals in the sequence and global coupling among sub-domains in the graph.

During fusion, we supply the concatenated sequence + structure embeddings as the queries and the surface embeddings as keys and values to a Transformer decoder. Cross-attention directly gauges the similarity and complementarity between sequence–backbone features and surface descriptors, aligning their shared patterns at driver-mutation sites. After several layers of cross-attention and feed-forward refinement, the decoder outputs a unified fusion vector that retains the unique information of each modality while achieving complementary optimization in global context—furnishing the downstream classifier with a highly discriminative multimodal representation.

### Model Training

#### Loss Function

The model produces a probability score indicating whether a single–point mutation is a driver. We compute the discrepancy between predictions and ground-truth labels with *Binary Cross-Entropy with Logits* (BCEWithLogitsLoss). Because this formulation fuses the Sigmoid activation with cross-entropy in a numerically stable way, it alleviates issues arising from extreme probability values and promotes smooth gradient flow, helping the network converge reliably.

### Optimizer

Training employs the Adam optimizer, whose adaptive learning-rate scheme blends first- and second-moment estimates of the gradients. This accelerates convergence while dampening the impact of gradient noise throughout training.

### Training and Validation Procedure

An epoch consists of a full pass over the training set followed by a single validation run. During training, batch losses are calculated with BCEWithLogitsLoss and back-propagated through Adam to update all learnable parameters. In validation, only forward inference is executed, and intermediate metrics—such as accuracy, precision/recall, and ROC-AUC—are recorded to monitor generalization and guide hyper-parameter tuning.

## Discussion

This study presents a novel end-to-end differentiable multimodal framework, MultiAlloDriver, which integrates protein surface information with sequence and structural features to achieve precise identification of cancer driver mutations. First, the ESM-2 protein language model was employed to capture long-range sequence dependencies, while GVP-GNNs enabled equivariant encoding of three-dimensional structural features. Concurrently, atomic-level surface geometry and physicochemical properties were reconstructed using the dMaSIF algorithm. A transformer-based architecture, leveraging self-attention and cross-attention mechanisms, facilitated global alignment and synergistic fusion of these multimodal features, ensuring complementary information flow across sequence, structure, and surface representations. Evaluated on a large-scale driver/passenger mutation dataset previously curated by our group, the model demonstrated robust performance, achieving an accuracy and precision of approximately 94% on the test set.

Its biological relevance was further validated through the identification of the F90S allosteric driver mutation in PTEN, where MultiAlloDriver successfully predicted its impact on dephosphorylation activity, underscoring the framework’s generalizability and mechanistic interpretability.

By pioneering the integration of protein surface information—a high-dimensional functional descriptor—into driver mutation prediction, MultiAlloDriver not only enhances the detection of complex allosteric sites but also provides new avenues to elucidate protein-protein and protein-ligand interaction mechanisms. Future efforts will focus on extending this framework to additional cancer-related genes and ligand-binding systems, refining model architectures and feature representation strategies. Such advancements are anticipated to yield computational tools with enhanced predictive power for targeted drug discovery and precision oncology.

## FUNDING

This study was supported by grants from the National Key R&D Program of China (NO.2023YFC3404700) the National Natural ScienceFoundation of China (No. 81925034).

## Notes

### Competing Interest Statement

The authors have declared no competing interest.

### Summary of Updates

The author's contribution has been revised

